# Functionalization of Gold Surfaces with Dithiobis(succinimidyl propionate) for Immobilization of Fetuin-A and Assessment of the Attachment and Proliferation of Osteoblast-like Cells

**DOI:** 10.64898/2026.05.05.722870

**Authors:** Alessandra Merlo, Jesper Medin, Andreas Dahlin, Kathryn Grandfield, Kyla N. Sask

## Abstract

Surface functionalization of biomaterials enables the immobilization of proteins and other molecules and can be utilized to direct the biological response to devices and implants. Fetuin-A is a blood plasma protein involved in numerous physiological processes, including the regulation of mineralization. Notably, many investigations of fetuin-A have explored its cellular interaction when in solution, but limited studies report the role of fetuin-A when used as a surface modifier. The present investigation explores the response elicited by fetuin-A on Saos-2 cells when it is immobilized on a model gold surface through the covalent reaction with dithiobis(succinimdyl propionate) (DSP). Comparative surface characterization using x-ray photoelectron spectroscopy (XPS), atomic force microscopy – infrared spectroscopy (AFM-IR) and surface plasmon resonance (SPR) confirmed the surface modifications but indicate partial inhomogeneity in the functionalizer surface coverage. The interaction of albumin and fetuin-A with the surface was quantified by radiolabeling, quartz crystal microbalance with dissipation (QCM-D) and SPR, demonstrating a higher mass of fetuin-A bound to the surface in comparison to serum albumin. Over 7 days, cells bound to the surfaces with immobilized fetuin-A showed significantly hindered proliferation of osteoblast-like cells compared to the positive control (fibronectin), presumably due to a decrease in cell metabolism. This study provides new insights into the role of fetuin-A in regulating Saos2 cell response and elucidates its potential use in combination with chemical functionalizers for biomedical applications requiring surface modification.

## 1. Introduction

Surface functionalization is a widely used strategy to improve or direct the biological response to materials that contact bodily fluids or tissues. While the choice of the biomaterial for specific applications is often constrained by its bulk mechanical properties, such as flexibility, elastic modulus, or permeability, the surface of the biomaterial can be selectively modified to tune its properties and thereby elicit specific effects on the surrounding tissue and aid in the restoration of native functionality. Modifications of either the mechanical^1^ or physical properties of the material surface, such as the presence of patterned, controlled^2,3^ or unordered, random surface topography^4^, have been shown to be effective in modulating the immunomodulatory response or increasing cellular proliferation and attachment.

The response to a biomaterial or implant can also be controlled through the variation of its chemical properties, commonly achieved via chemical functionalization. In this approach, the surface is modified to present specific moieties or compounds on the original substrate, deliberately selected to promote, for example, cellular adhesion and proliferation^5,6^, reduce bacterial interaction^7,8^, prevent protein adsorption^9^ or inhibit thrombus formation^10,11^, among other possible effects. Upon interaction of a biomaterial with physiological fluids, proteins rapidly bind to the surface through adsorption processes. Given its efficacy and simplicity, pre-adsorption of proteins or other nature-inspired molecules has been widely employed to pre-treat surfaces^12,13^. However, adsorption involves the formation of relatively weak non-covalent interactions, such as dipole interactions, hydrophobic and hydrogen bonds, and Van Der Waals forces^14^.

Treatments based on pre-adsorbed proteins have demonstrated limited efficacy, given the conformational changes and possible protein denaturation caused by adsorption^15^. In addition, upon interaction with biological fluids, pre-adsorbed molecules can be displaced by proteins with higher surface-affinity (Vroman effect)^16^, which further compromises the effectiveness of the engineered surface treatment. Therefore, to guarantee the stability and consistency of the surface functionalization over time, biological moieties are should preferably be bound to the surface through covalent bonds, which prevent the desorption or substitution of the pre-adsorbed molecule and maintain the desired surface characteristics over extended periods.

For bone-interfacing implants, various attempts have been made to engineer the surfaces with the goal of improving osseointegration and accelerating the restoration of the patient’s ability to perambulate. A wide range of moieties can be covalently immobilized onto implant surfaces to improve cell attachment^6^ and proliferation, or inhibit bacterial cell colonization^17,18^. The deposition of hydroxyapatite or calcium phosphate-based compounds is among the most commonly employed approaches^19,20^, owing to their compositional and structural similarity to the inorganic matrix of bone and their ability to recreate a bone-like microenvironment.

Fetuin-A is a molecule of growing interest in orthopedic and dental applications, given its established role in mineralization processes and its calcium/phosphate binding capabilities. Present in adult blood plasma at a concentration of approximately 0.4 mg/mL^21^, fetuin-A has been recognized as an antagonist of transforming growth factor-β (TGF-β) and various bone morphogenic proteins, which are crucial for bone formation and metabolism. The protein has also been shown to participate in cellular growth and proliferation processes^22^, particularly in certain tumorigenic cell lines^23,24^.

In a previous investigation by Oschatz et al^25^, fetuin-A was immobilized on a PLLA-co-PEG exploiting the 1-ethyl-3-(3-dimethylaminopropyl)carbodiimide / N-hydroxysuccinimide (EDC/NHS) coupling strategy. The authors observed increased hydroxyapatite (HA) deposition on fetuin-A functionalized surfaces following immersion in a supersaturated solution containing calcium and phosphate ions. The enhanced formation of HA on the functionalized surface suggested the potential of this protein as an effective surface modifier for bone-interfacing applications. However, while the effect of fetuin-A on various cell lines has been studied extensively in solution, relatively few studies have examined its role in cell attachment and proliferation when the protein is either physically adsorbed^26^ or covalently immobilized on a surface.

The present investigation explores the effect of fetuin-A on osteoblast-like cell response when the protein is chemically bound to a model gold surface through a self-assembled coupling agent, dithiobis(succinimdyl propionate) (DSP). Following surface functionalization, substrates were characterized using water contact angle measurements, X-ray photoelectron spectroscopy (XPS), atomic-force microscopy – infrared spectroscopy (AFM-IR) and surface plasmon resonance (SPR) to confirm the functionalization. Quantitative information regarding protein mass, binding kinetics and viscoelastic properties of the immobilized protein layer was obtained through comparative quartz crystal microbalance with dissipation (QCM-D), SPR and radiolabeling studies. Substrates were then evaluated for cellular attachment, proliferation and metabolic activity over a 7 day period to assess the potential role of surface-bound fetuin-A in promoting bone tissue regeneration.

## 2. Materials and Methods

### 2.1 Materials

Bovine serum albumin (BSA, 05470), bovine fetuin (BFet, F2379) and bovine fibronectin (BFn, F4792) were purchased from Sigma Aldrich and reconstituted as per manufacturer’s directions, and diluted to 0.2 mg/mL using phosphate buffered saline (PBS). Dithiobis(succinimidyl propionate) (DSP, D3669), dimethylsulfoxide (DMSO, 472301), McCoy Media 5A (M9309), penicillin/streptomycin (P4333), saponin (S4521) and formaldehyde 37% solution (F8775) were purchased from Sigma-Aldrich (Oakville, ON, CA).

### 2.2 Surface modification and protein immobilization

Gold-coated silicon wafers (100 nm gold with a 30-50 nm chromium adhesion layer) were purchased from Silicon Valley Microelectronics Inc (Buellton, CA) and diced into 0.5 x 0.5 cm pieces. All substrates were cleaned by immersion in a boiling solution of ultrapure water (Milli-Q), hydrogen peroxide and ammonium hydroxide (4:1:1 v/v) for 10 min, followed by rinsing three times in ultrapure water and nitrogen drying.

DSP modification was achieved by incubation of gold substrates in 20 mM DSP in DMSO for 2 h at room temperature to introduce NHS groups for protein immobilization, as previously described^27^. Following functionalization with DSP, the surfaces were rinsed with DMSO, washed in ultrapure water and then PBS and transferred to a 0.2 mg/mL protein solution in PBS for 3 h to immobilize proteins via reaction with the amino groups of lysine residues.

### 2.3 Water contact angle

Water contact angle measurements were performed using the sessile drop method with a Ramé-Hart NRL 100-00 goniometer (Mountain Lakes, NJ). Data are reported as average ± SD, with number of replicates (N) ≥12. Results were analyzed using the one-way Analysis of Variance (ANOVA) method. The significance level was defined at 0.05, with *p<0.05, **p<0.01, ***p<0.001 and **** p<0.0001.

### 2.4 X-ray photoelectron spectroscopy (XPS)

The XPS analyses were carried out with an AXIS Supra X-ray photoelectron spectrometer (Kratos Analytical, Manchester, UK) using a monochromatic Al Kα source (12 mA, 15 kV) on the bare gold substrates, the DSP-functionalized gold substrates (DSP) and then DSP-functionalized gold substrate with fetuin-A immobilization (DSP+BFet). Two replicates for each sample type were probed (N=2). The instrument work function was calibrated to give a binding energy (BE) of 83.96 eV for the Au 4f_7/2_ line for metallic gold and the spectrometer dispersion was adjusted to give a BE of 932.62 eV for the Cu 2p_3/2_ line of metallic copper.

The Kratos charge neutralizer system was used on all specimens. Survey scan analyses were carried out with an analysis area of 300 µm × 700 µm and a pass energy of 160 eV. Sample tilt of 0° was used to compare all samples, while sample tilt of 60° was considered for the DSP-FetA sample to be compared with the 0° tilt, to detect possible difference within chemical composition between surface and inner layers.

Spectra were analysed using CasaXPS software (version 2.3.26).

### 2.5 Atomic Force Microscopy – Infrared Spectroscopy (AFM-IR)

Dimension iconIR (Bruker, Billerica, MA, US) was used to perform Tapping AFM-IR measurements using a gold coated AFM-IR probe with a nominal spring constant of 40 N/m (PR-UM-TnIR-D). The fingerprint region spectrum was accessed by a tunable QCL (DRS Daylight solutions) equipped with the system. The tapping drive frequency operated at the second resonance mode f2 = 1605kHz while IR was detected at f1 = 260 kHz. Thus the laser was pulsed at the difference frequency of the two modes around 1345 kHz. Each IR spectrum represents a co-average of 3 at 4% laser power.

### 2.6 Surface Plasmon Resonance

SPR sensors (BioNavis, Tampere, Finland) were cleaned and surface-modified as described in Section 2.2., with 2 h submersion in a 20 mM DSP solution in DMSO. Samples were then rinsed in PBS, in milliQ water to eliminate saline residues, and dried with nitrogen. All measurements were collected on a BioNavis SPR Navi 220A instrument. A 670 nm laser diode was used to monitor total internal reflection (TIR) and SPR angle. SPR spectra analysis on ex-situ dry measurements was performed by Fresnel modeling according to the methodology previously reported^28^, and used to confirm the presence of DSP on the gold surface, and evaluate the thickness of the DSP layer.

For protein immobilization studies, in-situ measurements were performed following baseline stabilization for at least 10 min in PBS. The TIR and SPR angle were monitored with a 670 nm laser diode. The flow rate of PBS buffer (pH 7.4) was maintained constant at 10 μL/min, to conduct the observation under low flow conditions, but under analogous condition to the static protein immobilization used in Sections 2.2 for radiolabeling experiments and cell culture studies. The total *in situ* real-time protein adsorbed mass reported refers to the calculations on the 670 nm laser channel done according to previous work^28^, with maximum shift measured after 40 min (N≥3 replicates). Associations constants measured through SPR, intended as initial attachment kinetics calculated from the overall average response, were obtained using initial slope values. All measurements were done at room temperature. Following protein adsorption, 30 min rinses with PBS (pH 7.4) were performed, to rinse any loosely-bound protein.

### 2.7 Quartz-Crystal Microbalance with Dissipation

Quartz Crystal Microbalance with Dissipation (QCM-D) gold sensors (QuartPro, Stockholm, Sweden; 5MHz Cr/Au) were cleaned as described in Section 2.2. measurements were performed on a QSense Analyzer (Biolin Scientific, Gothenburg, Sweden), in PBS buffer (pH 7.4). Measurements were collected after baseline stabilization for at least 10 min. The flow rate used was 100 μL/min, to conduct the observation under low flow conditions, similar to the static protein immobilization used in Section 2.2. All the measurements were done at 25°C, with 30 min PBS rinses following protein adsorption, in order to rinse loosely-bound proteins. The mass adsorbed on the sensor was estimated through the Sauerbrey equation (Equation 1), using the fifth overtone and a *C* value of 17.7 ng/(cm^2^*Hz).

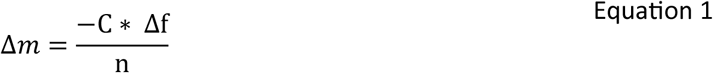

Association constants measured through QCM-D, intended as initial attachment kinetics calculated from the overall average response,were obtained using the Langmuir adsorption model (Equation 2). C(t) was assumed equal to C_0_ due to the constant liquid exchange,Γ(t) ≈0 for small t values, and Γsaturation=Γmax, given the limited degree of desorption during PBS rinses. The association constant can be calculated considering the value of 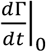 equal to the initial slope of the SPR in-situ real-time curves, as shown in previous work^29^. All modelled values are provided with a 95% confidence interval.

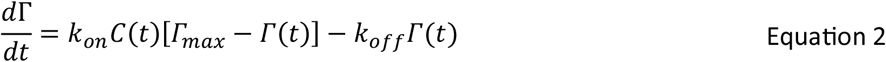

The total protein adsorbed mass reported, as well as the association constant, refer to the calculations from the 5^th^ overtone with maximum shift measured after 40 min (N≥2 replicates). To confirm the total amount adsorbed after 3 h adsorption and full surface coverage, longer runs of 160 min were conducted with the same parameters.

### 2.8 Radiolabelling with ^125^Iodine

Both BSA and BFet were labelled with Na^125^I using the iodine monochloride (ICl) method. The labelled protein was filtered with an AG 1-X4 anion exchange resin to eliminate free radioactive isotope. Labeled protein was precipitated in 20% trichloroacetic acid (TCA) and centrifuged for 5 min. The supernatant, containing free iodide ion (i.e. not bound to protein) was counted on a Wizard Automatic Gamma Counter (PerkinElmer, Shelton, CT, USA). Tests were conducted to determine residual free iodine, and levels were maintained below 3%. To prevent the binding of residual free iodine any section of the gold surface not fully covered by DSP, protein dilution and immobilization experiments were carried out using phosphate buffered saline containing nonradioactive sodium iodide (PBS–NaI) (3%)^30,31^, in a 1:9 mass ratio of labelled:unlabelled protein. BSA and BFet solutions in PBS-NaI buffer were prepared with final concentrations of 0.2 mg/mL, and then samples (N=4) were incubated for 3 h at room temperature in static conditions, as in Section 2.2. Following adsorption, the samples were rinsed three times with PBS, for 5 min each. The samples were lightly wiped to remove the residual adherent buffer, and the radioactivity was counted with a gamma counter. Radioactivity was converted to protein adsorption amounts based on corresponding solution counts. Following radioactivity counting, samples (N=4) were incubated in sodium dodecyl sulfate (SDS, 2% in water) overnight with shaking (250 rpm), to elute loosely adsorbed protein. The SDS was subsequently removed, and the samples were rinsed three times with PBS buffer for 5 min each. Finally, the samples were dried, and radioactivity measured through gamma radiation counts as previously described to obtain the residual surface coverage. Data are reported as average ± SD, and analyzed using the two-way Analysis of Variance (ANOVA) method. The significance level was defined at 0.05, with *p<0.05, **p<0.01, ***p<0.001 and **** p<0.0001.

### 2.9 Cell culture

Saos-2 osteosarcoma cells (ATCC®, HTB-85) were grown in McCoy’s 5A modified medium supplemented with 15% fetal bovine serum (FBS, Wisent Inc., St. Bruno, QC, CA) and 1% penicillin/streptomycin. All experiments were conducted with cell passage number ≤10. All samples were cleaned and functionalized as described in section 2.2 and then sterilized with 30 min exposure to UV light on both sides of the substrates. To seed cells on the substrates, cells were treated with 0.25% Trypsin-EDTA for 6 min at 37°C. Immediately afterwards, an equal amount of non-supplemented media was added, and cells were centrifuged for 5 min at 1000 x g, the eluted solution removed, and the cell pellet resuspended in non-supplemented media. Cells were then seeded in non-supplemented media and incubated for 24 h at 37 °C, 5% CO_2_. After 24 h, non-supplemented media was substituted for McCoy’s 5A modified medium supplemented with 15% FBS and 1% PS, and culture media exchanged every two days.

### 2.10 Fluorescence Microscopy

Before staining, substrates were washed twice with PBS to eliminate non-adherent cells present on the surface. Then substrates were flash permeabilized with a 0.1% saponin solution for around 10-15 seconds, then fixed with a 4% formaldehyde solution for 7 min at 4°C. Samples were washed twice with PBS. A 2% BSA blocking solution was added for 1 h at 37°C. Substrates were washed twice with PBS, and a staining solution containing nucleic stain (Hoechst 33342, Life Technologies Inc., CA, USA) and phalloidin stain (phalloidin-Alexa fluorophore 488) in PBS (3:1:3000 v/v) was added for 2 h at room temperature. Samples were washed with PBS and then mounted face-down with Fluoromount-G Mounting Medium (00-4958-02, Life Technologies Inc., CA, USA) on glass coverslips. Cell attachment and proliferation were assessed using immunostaining on days 1, 3, and 7. Cells were imaged on a Nikon A1R Inverted Confocal with a 20x air objective lens (NA 0.75) on day 1, using the large image function embedded in the NIS Viewer software to image a larger surface area. On day 3 and day 7, a 10x air objective lens, (NA 0.45) was used.

### 2.11 Image Analysis

#### 2.11.1 Cell Nuclei Count

The software CellProfiler was employed for the cell nuclei count on day 3,, with the following workflow: the DAPI channel of collected images was thresholded in 2 classes using the Ostu method, and nuclei identified through the Identify Primary Objects functions using the same thresholding method discarding objects <30 and >350 px outside of the size range of single nuclei. For day 7 images, the confluency over the monolayer required a first segmentation stage using the software Ilastik to separate nuclei from background through a Pixel Classification process. Output masks were then opened in Fiji software, thresholded with the Ostu method, as done in CellProlifer, to create a binary mask, and then the number of objects counted through the Analyze Particles function with the same thresholds mentioned above. For each condition, ≥3 images on random areas of 3 replicates, per condition, were averaged. Each of these averages was then averaged with 3 replicates per sample (Average), and then statistically compared with other substrates. Results are presented as Average ± SD. Statistical differences were measured using an independent two-way ANOVA with Tukey’s HSD post-hoc test in GraphPad Software (San Diego, CA, USA). The significance level was defined at 0.05, with *p<0.05, **p<0.01, ***p<0.001 and **** p<0.0001.

#### 2.11.2 Cell Surface Coverage

The total surface coverage (% of total area) of the cells was evaluated through the Ilastik software using the Pixel Classification method. 3 random images of the dataset, FITC channel, were used to define two classes, cell area and background, and train the model, which was then applied to the full dataset. Exported segmentations were then loaded in Fiji software, images converted into binary masks, and pixel for the two categories counted to define surface coverage. For each condition, ≥3 random regions of interest of the single sample were averaged. Each of these averages was then averaged with 3 replicates per sample (Average), and then statistically compared with other substrates. Results are presented as Average ± SD. Statistical differences were measured using a two-way independent ANOVA with Tukey’s HSD post-hoc test in GraphPad Software (San Diego, CA, USA). The significance level was defined at 0.05, with *p<0.05, **p<0.01, ***p<0.001 and **** p<0.0001.

### 2.12 Cell metabolism

Cell metabolism was assessed with an alamarBlue^TM^ assay (Life Technologies Inc., Carlsbad, CA, USA) on days 1, 3 and 7, with 6 replicates for each condition,and two repeats (N=12). An Infinite M200Pro plate reader (Tecan Group Ltd., Switzerland) was used to measure the fluorescence intensity values at 540 nm excitation and 580 nm emission wavelengths. The values of the two repeats were normalized over the average of day 1 - control value (polystyrene plate) of the corresponding run, then pooled together.

Statistical differences were measured using a two-way ANOVA with Tukey’s HSD post-hoc test in GraphPad Software (San Diego, CA, USA). The significance level was defined at 0.05, with *p<0.05, **p<0.01, ***p<0.001 and **** p<0.0001.

## 3. Results and Discussion

### 3.1 Surface Characterization

#### 3.1.1 Water Contact Angle

To verify the presence and homogeneity of the DSP surface modification, wettability analysis was performed using sessile-drop water contact angle (WCA) measurements. The results from the WCA measurements are presented in Figure 1. DSP functionalized surfaces presented a contact angle around 40°, similar to the contact angle measured on bare gold^32,33^. These values are consistent with previous studies using the same protocol for DSP chemisorption to gold^27^. A difference was found between DSP, without any protein bound, and all substrates where DSP was used to immobilize proteins, with each demonstrating an increase in hydrophobicity. This increase in surface hydrophobicity following protein adsorption may suggest a particular orientation of the proteins on the surface. No significant differences between the wettability of the protein-modified samples was found which could be related to the mechanism of protein immobilization, where the proteins bind to the NHS ester of DSP through their lysine residues, which in turn may limit the 3D rearrangement of the protein upon surface interaction.

**Figure 1.**
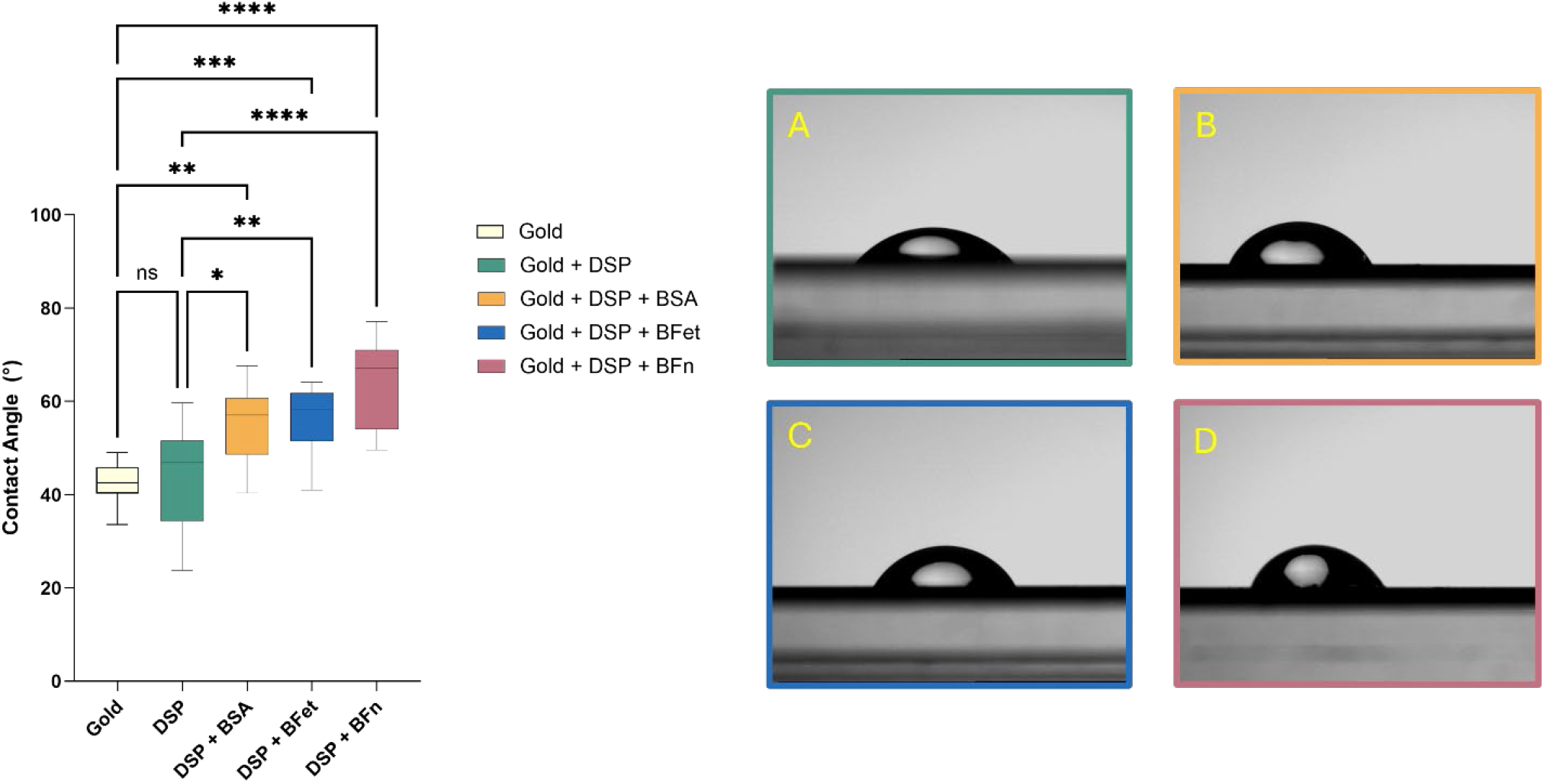
Water contact angle measurements obtained through the sessile drop method. Data are presented as averages ± SD, N ≥ 9. Statistical significance was evaluated through one-way ANOVA with * indicating p < 0.05; **, p < 0.01; ***, p <0.001, ****, p<0.0001. On the right side, representative images of droplets on a gold surface modified with DSP (A), gold surface modified with DSP and with BSA (B), BFet (C) or BFn (D).

#### 3.1.2 X-ray Photoelectron Spectroscopy

The presence of DSP on the surface, and its binding of fetuin-A, was confirmed through both x-ray photoelectron spectroscopy (XPS), with results summarised in Table 1 and Table 1S and 2S, atomic force microscopy – nano infrared (AFM-IR), presented in Figure 2, and surface plasmon resonance (SPR), presented in Figure 2S.

**Table 1.**
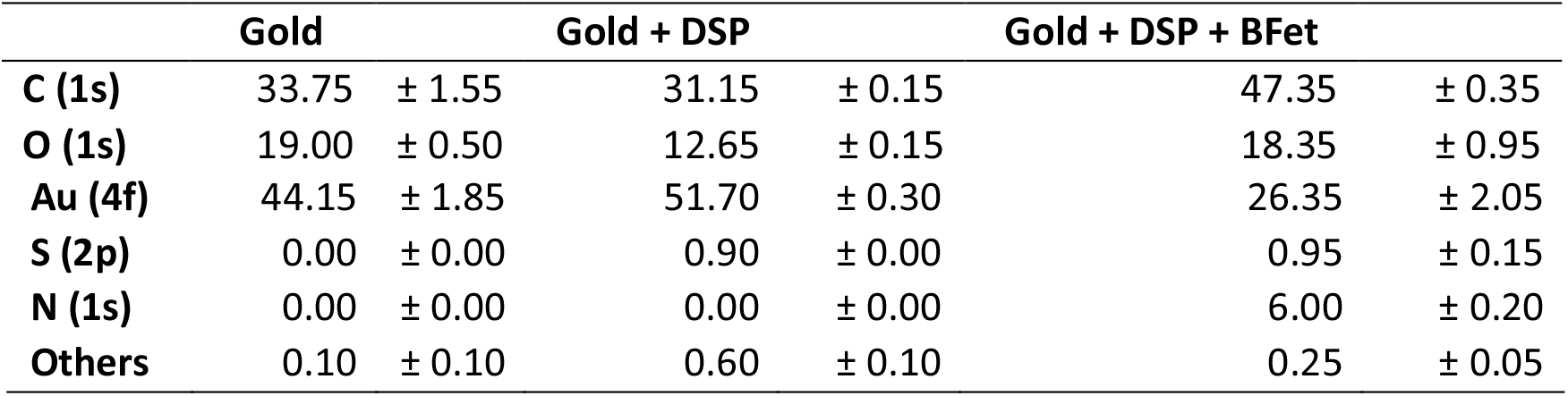
XPS data (at.%) at take-off angles of 90° comparing the surface composition of the DSP functionalized and protein-immobilized samples. Data are presented as mean ± range, N=2.

**Figure 2.**
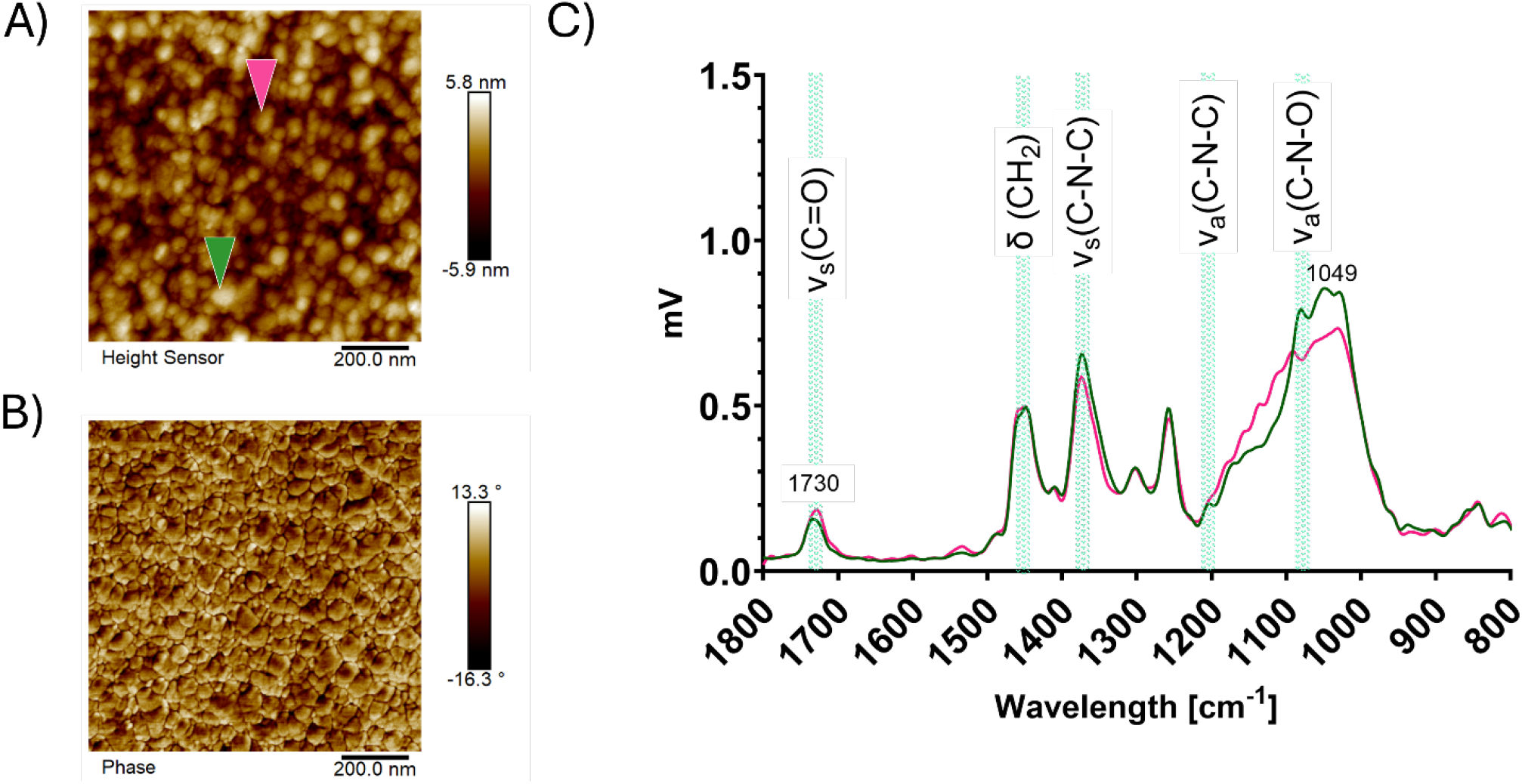
Atomic Force Microscopy - Infrared Spectroscopy analysis. A) displays the morphology of the Gold+DSP sample in tapping mode, and B) the phase lag in cantilever oscillation for Gold+DSP. C) IR spectra obtained for Gold+DSP The areas of the sample where the spectrum was obtained are shown with a downward arrow in the same color of the corresponding spectrum. The IR spectrum for Gold+DS confirmed the presence of functionalization on the gold surface.

Table 1 presents the average atomic percent of two different replicates for each of the three samples: the bare gold surface (Gold), gold coated with the surface functionalizer (Gold+DSP), and the functionalized gold surface with immobilized fetuin-A (Gold+DSP+BFet), representative of surfaces with immobilized proteins. The 2p sulfur peak from the DSP functionalizer, was absent on the bare gold surface, but appears with an atomic percentage of 0.9 ± 0.0 on the DSP surface, confirming the binding through thiolation and the formation of a self-assembled layer. Its varying distribution throughout the sample is evident by similar levels of heterogeneity on different replicates. The sulphur content increases to 1.0 ± 0.2 when protein is immobilized, as expected by the amino acid composition of the protein^34^. The 1s nitrogen peak is not detected for the functionalized surface Gold+DSP (Table 1), indicating either an amount of NHS too low for detection, possibly due to partial hydrolysis of DSP. The 1s nitrogen peak is strongly present with a 6.0 ± 0.3 atomic percentage for the substrates with immobilized protein. The detection of a relevant nitrogen peak for the Gold+DSP+BFet substrate, combined with an increase in C and O counts, confirms the presence and binding of the protein on the sample. The presence of trace contaminants on the surface, not related to functionalization or proteins, is indicated in the Others row, and further details on these are provided in Table 1S. Confirmation of the successful surface modification is also provided by the comparison of atomic concentrations collected at 0° and 60° degrees tilt angles (Table S2). The surface concentration of gold decreases for increasing tilt angle (higher surface sensitivity), while all signals related to protein or DSP remain constant or increase.

The binding energies assignments, and surface amounts detected, are in line with previous observations^35,36^. The competitive reaction between aminolysis and hydrolysis of DSP has been previously documented^37^, and reports have shown the NHS-ester group of the molecule to be prone to hydrolysis even when exposed to humidity in air, non-anhydrous solvents or throughout storage^38^. Such short and long-term instability of the molecule could explain why the nitrogen peak was not detected for the DSP-coated samples DSP was bound to the surface, as shown by the S peak.

#### 3.1.3 Atomic Force Microscopy – Infrared Spectroscopy

The successful binding of DSP and subsequent protein immobilization was also confirmed by atomic force microscopy-infrared spectroscopy (AFM-IR), as presented in Figure 2. Figure 2A shows the morphology of Gold+DSP samples, with a peak-to-valley roughness around 12 nm. The measured phase component, presented in Figure 2B for Gold+DSP, indicates an absolute phase angle of around 15°. The measured phase relates to the mechanical response of the underlying substrates, providing insight into the variation of stiffness or viscoelastic properties of the sample. The value detected, and the overall homogenous distribution on the area measured, suggests a stiff substrate. The results align with the expected measurements for the stiff gold-coated silicon wafer used for the analysis, where the presence of a monolayer of DSP would negligibly impact the measured stiffness of the overall substrate. The IR spectrum collected on the sample Gold+DSP (Figure 2C) in two different spots displays analogous peaks to those found in literature to confirm the presence of DSP on gold surfaces. In their investigation of a DSP monolayer chemisorbed on gold, Lim et al ^37^ measured characteristic peaks of the molecule: the N-C-O signal of the succinimide group at 1074 cm^-1^, the combined asymmetric and symmetric C-N-C stretch of the NHS at 1215 cm ^-1^ and 1378 cm^-1^, respectively, the methylene scissors deformation at 1464 cm^-1^ and finally the asymmetric carbonyl stretch of NHS at 1748 cm^-1^. Similar peaks, while slightly shifted, were detected in other works characterizing the DSP layer through FTIR^36,39^. In the present study, the peak of the C=O stretch of NHS is likely appearing at 1730 cm^-1^, and all other peaks, shown along the coloured spectra, match those found in the literature. As shown by Wagner et al.^40^ on a similar NHS ester self-assembled monolayer, the 1750 cm^-1^ peak can display as a broader band deriving from two unresolved bands at lower frequency (1744 cm^-1^ and 1729 cm^-1^, respectively). The hydrolysis of the NHS group leads to the disappearance of the above-mentioned absorption peaks in such regions, confirming the presence of active NHS. In addition, AFM-IR mapping results for the protein-immobilized samples provided further insight on the protein coverage and distribution (Figure 1S).

#### 3.1.4 Surface Plasmon Resonance

Surface Plasmon Resonance (SPR) measurements performed on the sample after 2 h submersion in DSP solution in DMSO show a change in SPR angle, supporting the presence of an additional layer on top of the gold sensor surface (Figure 2S, I). The thickness of the layer, evaluated through Fresnel modeling, indicates a deposited DSP layer with an average thickness of around 0.27 ± 0.05 nm. This value is less than the theoretical value of a DSP monolayer of 1.2 nm, calculated considering a half chain length for the molecule to account for thiolation. Two hypotheses could explain this difference: the first is that the layer of deposited DSP is not a homogeneous monolayer. The second is that the DSP molecule is not arranged vertically, exposing the sulfur to the gold substrate and the NHS group to the solution, but in an angled orientation with respect to the gold surface, thus decreasing the coverage density of the molecule, the thickness of the layer chemisorbed, and possibly limiting the exposure and availability for protein to bind to the NHS groups. Previous reports indicating the monolayer structure and orientation are dependent on alkane chain length and SAM preparation steps support the latter hypothesis^41^.

Our observations suggest that the generally accepted method previously adopted in multiple publications, where FTIR is used for verification of the formation of a complete DSP monolayer, is not sufficient for evaluating the homogeneity of the layer nor the full surface coverage from protein solution, and that further characterization of the system should be pursued to obtain additional results. It is possible that subsequent protein binding includes a combination of multiple phenomena at the interface: the protein forms covalent bonds with the chemisorbed layer, but also adsorbs on the bare gold surface in non-homogenously covered sections, possibly also interacting with partially hydrolyzed layers.

### 3.2 Protein characterization

The immobilization of BFet on the surface was studied in comparison to BSA. While the two proteins’ amino acid sequences are quite dissimilar, and they perform different physiological functions, serum albumin and fetuin-A are comparable in both molecular weight and globular structure in solution: 66 kDa molecular weight ^42^ and 3.51 nm hydrodynamic radius ^43^ for BSA, compared to 60 kDa ^34^ and 4.2 nm ^44^ for fetuin-A.

The quantification of protein bound to the DSP-coated gold substrates was obtained through the use of three techniques: SPR, radiolabeling and quartz crystal microbalance with dissipation (QCM-D). It is important to note that the mass coverages measured by SPR and radiolabeling correspond to the net protein coverage, whereas the mass measured by QCM-D is greater than the total mass of protein bound since it includes water associated with the protein, also providing insight on the hydration and packing of the protein layer at the interface. The comparison of results from these three techniques provided insight on various other properties of the immobilized layers: binding kinetics, water entrapment, and remaining protein levels after PBS rinses or treatment with SDS surfactant to elute loosely bound protein.

#### 3.2.1 Surface Plasmon Resonance and Quart-Crystal Microbalance with Dissipation

Figure 3A presents a comparison of the amount of protein mass bound to the surface measured through SPR, in dotted lines, and QCM-D, in solid lines. The immobilized protein mass measured by SPR after 160 min of BSA injection plateaus at a value of 133 ng/cm^2^ (dotted yellow line). BFet (dotted blue line) displayed a total amount of protein bound equal to 159 ng/cm^2^. Considering the similar molecular mass of BSA and BFet, the higher total mass immobilized for fetuin-A indicates a higher amount of BFet molecules immobilized on DSP-coated substrates, compared to BSA, from equally concentrated protein solutions.

**Figure 3.**
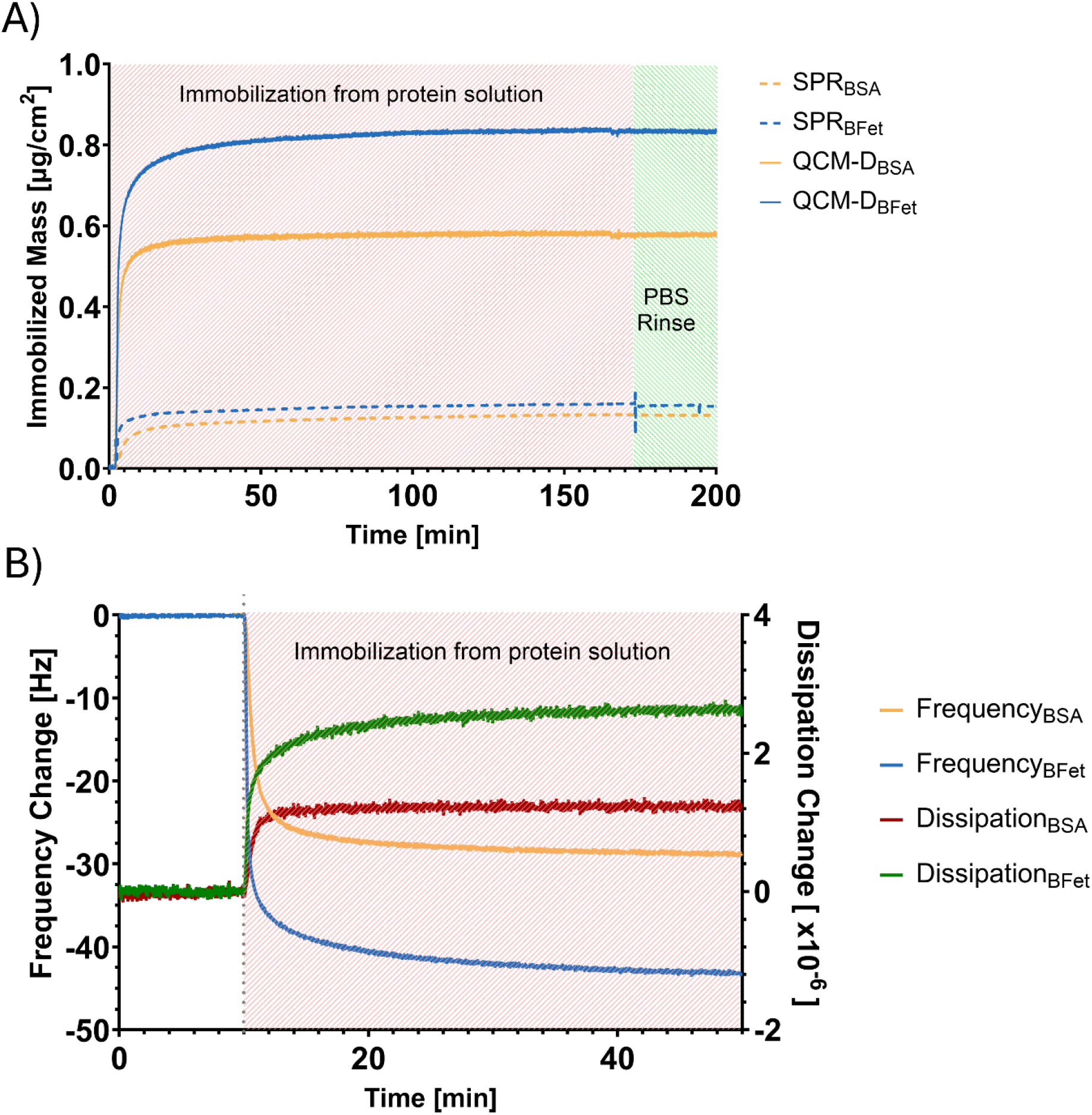
(A) Comparison of mass adsorbed to the Gold+DSP surface up to saturation detected through quartz crystal microbalance with dissipation (QCM-D) and surface plasmon resonance (SPR). Solid lines represent QCM-D, 5th overtone, indicating the hydrated weight of the protein immobilized through DSP; dotted lines represent the net weight of adsorbed protein through SPR. BSA is presented in yellow, BFet in blue. Both techniques demonstrate a higher amount of fetuin-A bound compared to the control protein on the functionalized surface, with the higher association constant attesting to a stronger bond between functionalized surface and BFet. (B) Comparison of frequency and dissipation changes observed in QCM-D upon immobilization of BSA and BFet on DSP-coated gold surfaces. The grey, dotted line indicates the injection of the protein solution (0.2 mg/mL in PBS) after baseline stabilization for 10 mins in PBS. The higher measured dissipation for BFet is to be related to less compact and more hydrated protein layer compared to BSA.

Previous results that measured the adsorption of these molecules on bare gold surfaces through SPR and QCM-D^29^ also highlighted a higher amount of BFet bound to gold surfaces compared to BSA. The initial slopes, indicative of change in mass over area, are 0.08±0.02 µg cm^-2^ min^-1^ for BSA and 0.16±0.04 µg cm^-2^ min^-1^, for BFet, indicating a quicker binding of BFet to DSP-functionalized surface with respect to BSA, as previously observed on bare gold substrates ^29^.

The difference in mass bound between BSA and BFet previously observed for proteins adsorbed directly to gold^29^ is, on the other hand, smaller when the surfaces are chemically functionalized with DSP: after 40 minutes of injection, BSA mass bound to the surface increases by about 12% when the protein is immobilized, while BFet mass bound decreased by about 14% on DSP, when compared to adsorption to bare gold surfaces^29^. If proteins are able to bind to DSP covalently through lysine residues of the proteins, this may result in limited structural rearrangement upon surface interaction. This hypothesis is further reinforced by the absence of a linear region in the SPR curve for BFet immobilization, which was present when the protein adsorbed on bare gold surfaces^29^ (Figure 3S). The difference between the two trends suggests limited re-arrangement upon immobilization on the DSP-coated substrate. A restriction in 3D rearrangement of the proteins may, in turn, vary the protein surface coverage, explaining the opposite trend in surface coverage observed for BSA and BFet when adsorbed on bare gold surface or covalently-immobilized through DSP. In addition, the intrinsic differences between the two proteins and their conformation in solution, as well as the number and accessibility of lysine residues, could lead to a change in packing density of the protein on the surface, explaining the difference in net protein mass bound to DSP-coated surfaces between BSA and BFet, as well as justifying the reduced difference in protein mass bound to the surface with respect to the bare gold case^29^.

For both proteins, rinsing with PBS (pH 7.4) for 20 min, under flow to remove loosely-bound proteins, led to a very limited reduction in immobilized mass, indicating stable binding of both proteins to the functionalized substrate. The amount of protein immobilized on the substrates was confirmed with complementary dry ex-situ SPR measurements: following 3 h submersion of the gold-coated samples in 0.2 mg/mL protein solution, PBS washing and nitrogen drying, similar differences in protein mass bound were identified: BFet bound in higher amount to the surface than BSA (272.2 ±29.9 ng/cm^2^ for the former, 179.9±6.8 ng/cm^2^ for the latter). The resulting layer thickness and calculated mass adsorption are presented in Figure 1S (II and III).

Representative curves for frequency and dissipation obtained with QCM-D for both proteins, over 40 min injection, are shown in Figure 3B. The mass adsorbed measured through QCM-D at plateau conditions after 40 min (Figure 3B, solid lines) is notably different between the two proteins (758 ng/cm^2^ for BFet and 507 ng/cm^2^ for BSA, respectively), confirming higher BFet immobilization and the formation of a more viscoelastic layer, as highlighted from the splitting of the overtones (Figure 4S). On the other hand, bovine serum albumin presented a more marked splitting of overtones with respect to the case of adsorption on gold (Figure 4S, A), indicating a layer with higher viscoelasticity when the protein is immobilized through DSP. The difference in layer rigidity observed for BSA comparing the adsorption on bare gold and immobilization through DSP confirms the difference in protein layer characteristics and packing based on the binding mechanism.

The initial slopes, measured in QCM-D, are 0.56 ± 0.07 µg cm^−2^ min^−1^ for BSA and 2.07 ± 0.08 µg cm^−2^ min^−1^ for BFet, confirming quicker binding of BFet to the substrates. Over 40 min injection, QCM-D results report a lower amount of protein mass bound to the substrate in the case of DSP-mediated immobilization for both proteins: a decrease of 13% in surface coverage for BFet, and an 6% decrease for BSA with respect to the adsorption case on model gold surfaces^29^. On the contrary, after 3 h injection, QCM-D results report higher amount of protein mass bound to the substrates in the case of DSP functionalization: +25% for BSA, and +11% for BFet. For both BSA and BFet, in all QCM-D experiments run in the present investigation, ΔD/Δf < 10^−7^, satisfying the condition of the Sauerbrey equation. Subsequently, the difference between hydrated mass (from QCM-D) and net mass (from SPR) of protein immobilized corresponds to the additional mass of water added to the film. The data indicate similar hydration of the protein layer for both proteins (76% hydration for BSA, 78% for BFet), which differs from the bare gold case^29^. In presence of DSP-functionalization, BFet is immobilized on the surface in higher amounts than BSA, but the average number of water molecules associated per protein is similar.

A buffer-assisted rinse under uniform flow conditions using PBS, to eliminate any loosely bound protein from the surface, resulted in a negligible reduction in the amount of protein retained on the surface.

Association constants, as measured using the method described in Section 2.7, were 17.3*10^3^±0.2*10^3^ min^-1^ M^-1^ for BSA and 51.6*10^3^±2.9*10^3^ min^-1^ M^-1^ for BFet. The association constants of the two proteins measured for DSP-functionalized substrate are similar to those observed previously for bare gold model surfaces^29^.

#### 3.2.2 Radiolabeling

Protein quantification obtained through radiolabelling (Figure 4) demonstrates similar immobilized amounts for BFet compared to BSA, with no statistical significance detected. The discrepancy in amount of protein immobilization between radiolabeling and SPR/QCM-D (Figure 1S, IV) is hypothesized to be related with the inherent differences in the measurement: under static condition for radiolabeling, and under flow condition for both SPR and QCM-D. Shaking and longer elution with a surfactant (SDS) resulted in a notable decrease of residual protein immobilized on the surface. Both the lack of difference between the two protein amounts upon immobilization, as well as the partial elution of protein by SDS, suggests that there may be some surface sites not fully covered with DSP and a small amount of protein physically adsorbed, and thus easier to elute. It is possible that the presence of NaI in the buffer to prevent adsorption of residual free iodine directly to the uncoated gold surface may compete somewhat with the thiol-gold bond, in turn decreasing the DSP functionalization of the surface, and reducing its homogeneity and efficiency of protein immobilization.

**Figure 4.**
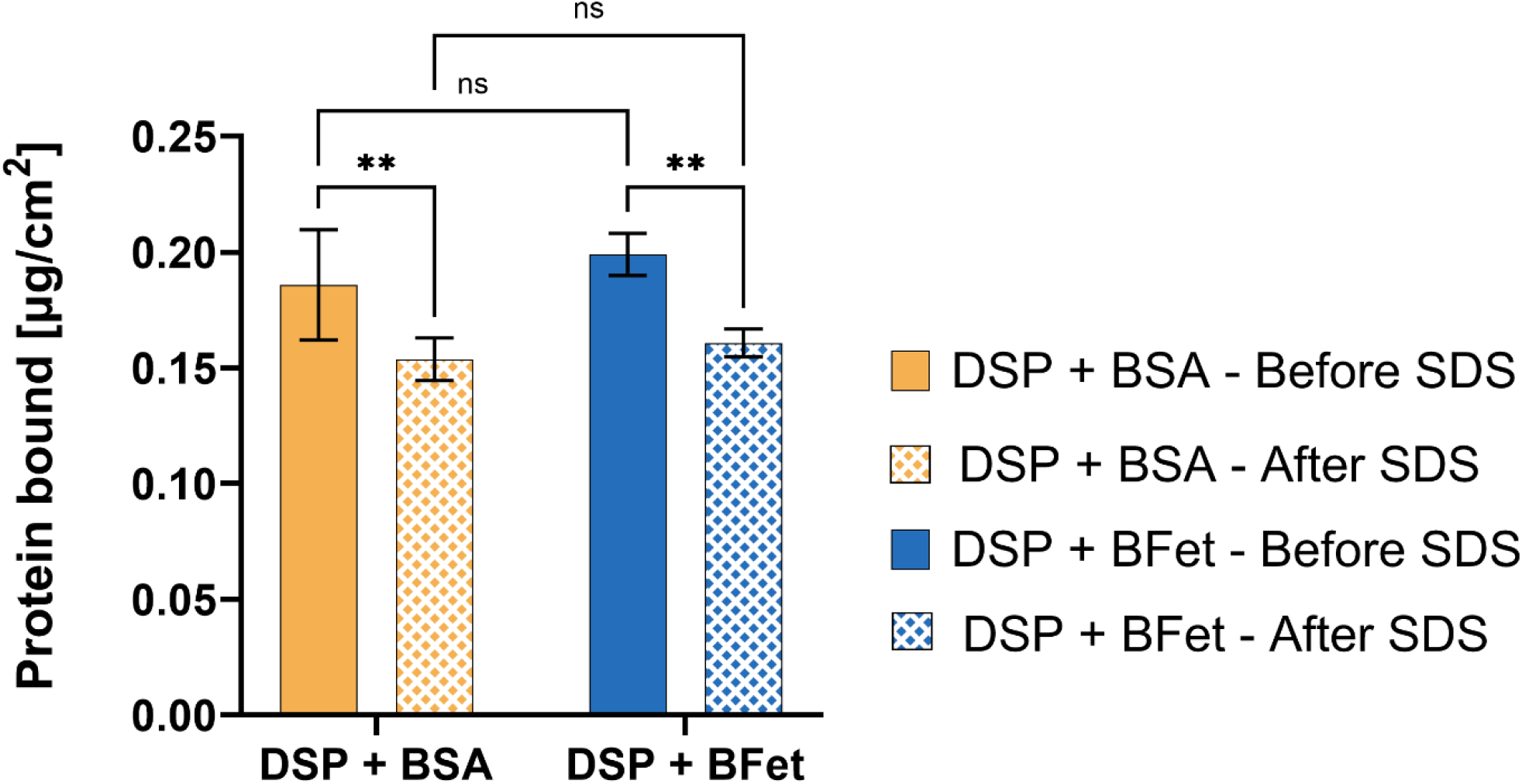
Protein surface density as measured by protein radiolabelling with ^125^I after 3 h of static incubation (before SDS) and following overnight elution in 2% SDS (after SDS, textured columns). Data are presented as averages ± SD, N=4. Statistical significance was evaluated through two-way ANOVA with ** indicating p < 0.01. BFet binds in similar amounts as BSA to gold surfaces, with a total adsorbed mass of around 0.2 µg/cm^2^ after 3 h in 0.2 mg/mL protein solution. Compared to previous findings, in the presence of DSP, there is a higher retainment of BFet following surfactant rinsing than for bare gold, confirming the stronger covalent binding.

Compared to the previous studies involving protein adsorption directly on gold^29^, it is evident that while BSA bound to surfaces in similar amounts in both circumstances, DSP-modified surfaces retained a higher amount of BFet. Such findings can be attributed to the creation of a stronger bond between the surface and the protein through the DSP functionalizer. These protein radiolabelling results confirm this since SDS can disrupt non-covalent bonds, but is ineffective when the proteins are immobilized through the DSP molecules.

### 3.3 Cellular Response

Fluorescence microscopy was employed to probe any inherent effects of immobilized fetuin-A on the proliferation of osteoblast-like cells. The results were interpreted based on observed variations in cellular metabolism detected through a fluorescence assay over 7 days. To prevent the immobilization of other serum proteins to any free NHS moieties still present or other sites available on the surface, osteoblast-like cells were first seeded in non-supplemented media^45^. Previous studies confirmed that there were no detrimental effects on cellular metabolism from the use of non-supplemented media^45^.

The ability of BFet to significantly affect cell adhesion, attachment and proliferation was compared to two other proteins, whose influence on cellular response have been more widely investigated. Fibronectin (BFn) is a plasma protein presenting the arginine-glycine-aspartic acid (RGD) sequence along its chain, recognized by the integrins of cells for the formation of stable binding sites to the extracellular matrix^46–49^. Given its cell-adhesive properties, BFn was selected as the positive control for the investigation. Albumin (BSA) was used as a negative control, although it lacks an RGD sequence, since it is the most abundant protein in plasma, and can act as a surface passivating agent.

#### 3.3.1 Cell Nuclei Count

No data could be collected after fixation of the DSP surfaces on day 1. Upon fixation and staining, the DSP surfaces did not present any attached cells on either substrate when observed under the microscope, regardless of the presence or absence of proteins. Only the remnants of stained cell actin filaments remained on the surface (Figure 5S), thus making any further evaluation of cell count and morphology at this time point impossible. It was not possible to determine the cause of cell detachment and wash-out, but it is hypothesized to be related to the lack of supplements in cell media, given the successful fixation at later time points.

**Figure 5.**
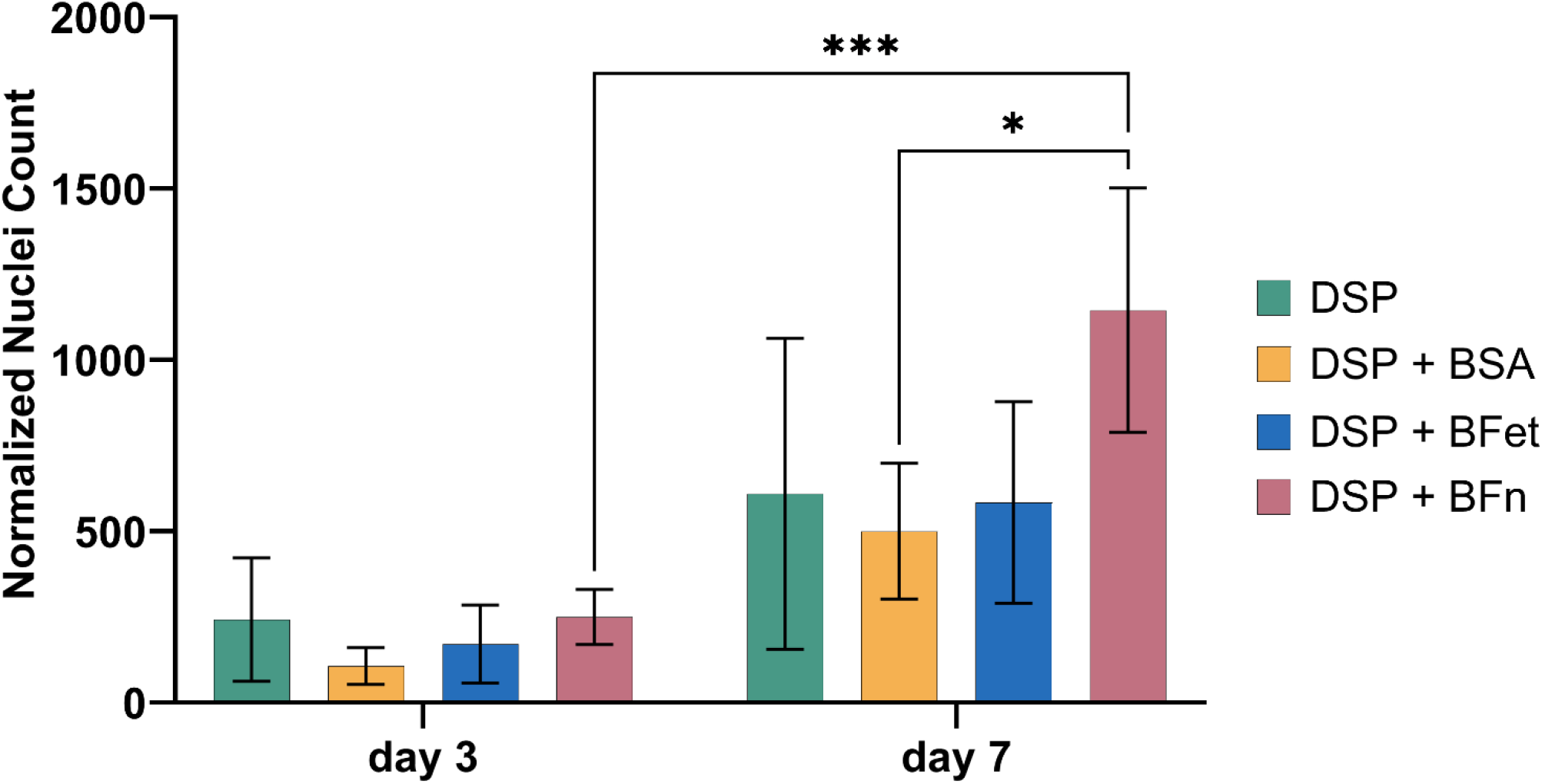
Total cell nuclei number on either the DSP-modified gold surface (DSP) or DSP-modified + protein surface at different time points (day 3, 7). Data are presented as Averages ± SD. Statistical significance was evaluated through two-way ANOVA with * indicating p < 0.05, ***, p<0.001. No statistical difference was found between BFet and other proteins at either of the time points.

On day 3, no statistically significant difference was found among the different samples (Figure 5). On day 7, DSP+BFn performed significantly better than DSP+BSA in promoting cell proliferation. DSP+BFet did not present statistically significant differences compared to neither the BFn positive control nor the BSA negative control. As previously observed, then, fetuin-A does not seem to have any inherent capability in promoting cell proliferation over time.

#### 3.3.2 Cellular Surface Coverage of Substrates

The nuclei count results are confirmed by the % surface coverage on the difference substrates (Figure 6). Over 3 and 7 days, no statistically significant difference was detected among the samples. In contrast to what was previously observed through physical adsorption to gold^29^, cell proliferation did not reach confluency over 7 days on most surfaces, with and without protein immobilization, indicating an overall decrease in cell proliferation in the presence of protein covalently-bound through DSP.

**Figure 6.**
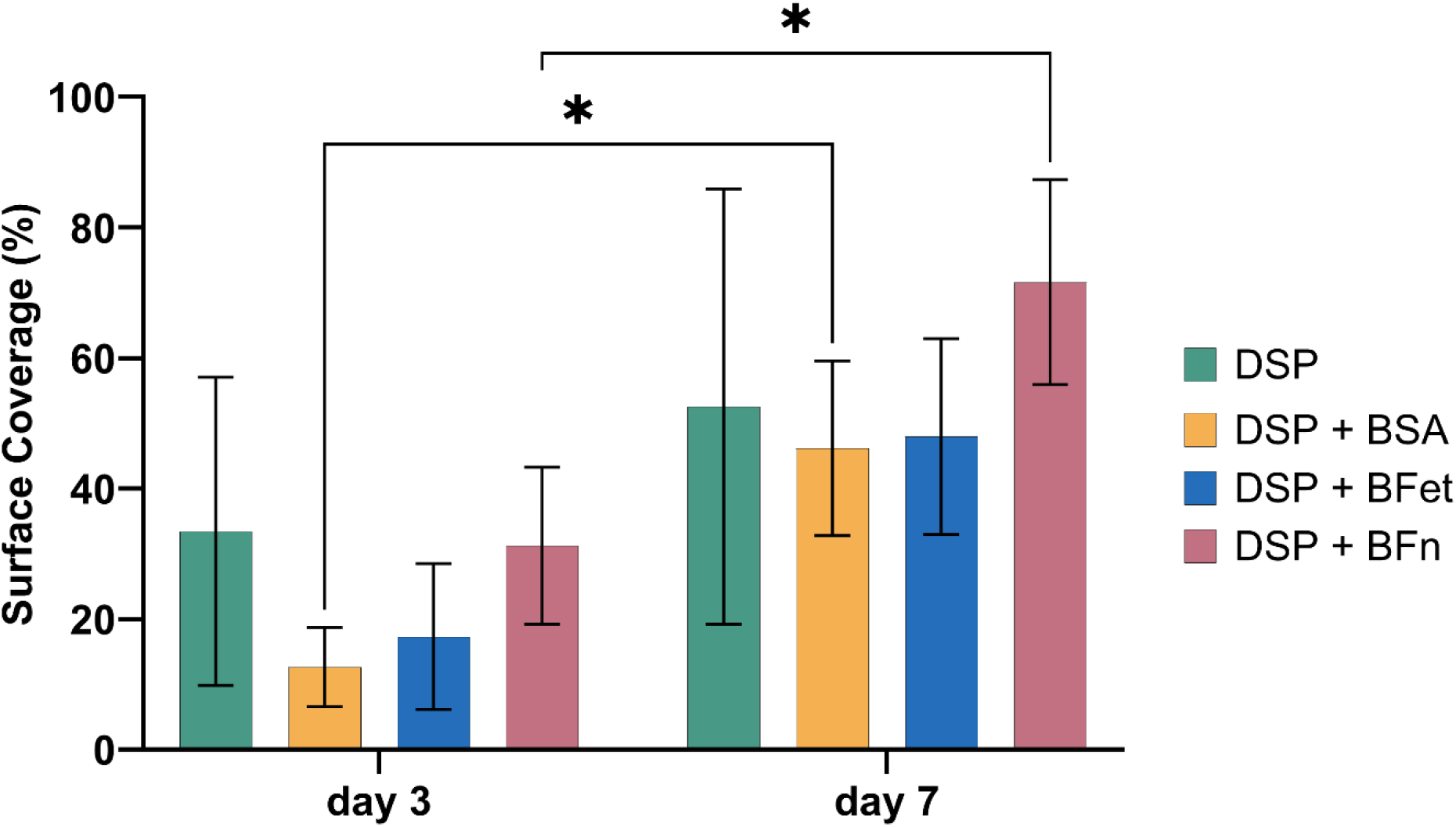
Percentage area of the sample covered by cell bodies on either the DSP-modified gold surface or DSP-modified + protein, at two different time points (day 3, 7). Data are presented as Averages ± SD. Statistical significance was evaluated through two-way ANOVA with * indicating p < 0.05. . No statistical difference was found between BFet and other proteins on either of the time points, similarly to what was observed in the nuclei count.

Such findings seem to confirm the hypothesis of a difference in protein rearrangement between the physically adsorbed and covalently-bound conditions. In the former, the protein interacts with the substrates through secondary, weaker bonds, such as hydrophobic or hydrogen bonds. In the latter, the covalent binding occurs through the substitution of the NHS ester moiety with protein amine groups either on the N-terminus or on side chains along the protein. As such, the 3D re-arrangement of the covalently-bound protein is likely limited, and the protein moieties generally exposed or utilized for cell interactions may be hindered from achieving their function. In addition, the formation of a monolayer of DSP molecules on the surface, each presenting a reactive moiety, can affect the packing density of the protein on the surface, and in turn, may induce some steric hindrance limiting the reactivity of the immobilized proteins.

#### 3.3.3 Cellular Metabolic Activity

A metabolic activity assay, AlamarBlue^TM^ HS (AB), was employed to determine any correlation in attachment and proliferation to the metabolic activity of the cells over 7 days. The results of the assay are presented in Figure 7.

**Figure 7.**
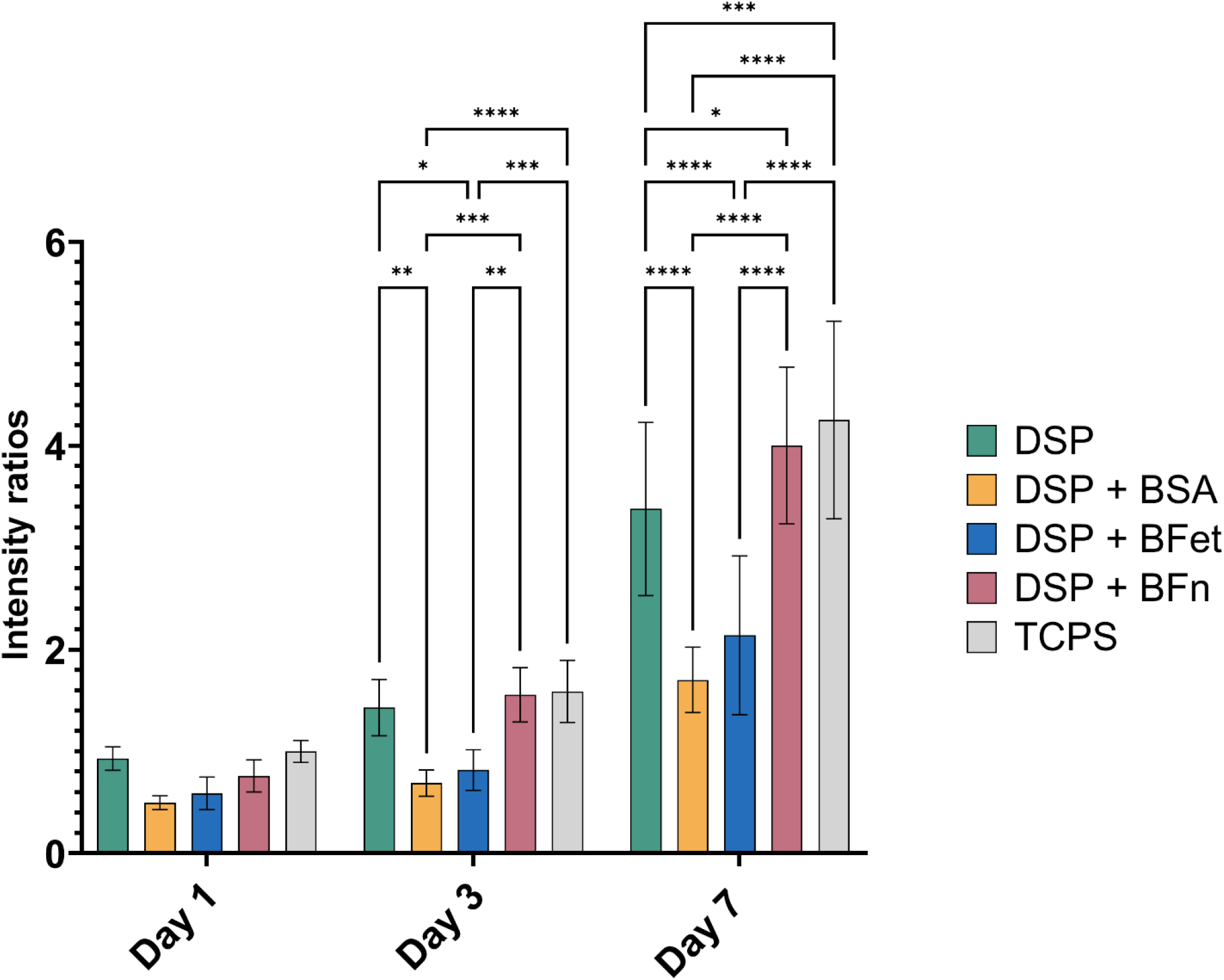
Fluorescence intensity values representing overall cell metabolism for cells grown on either DSP-coated gold or DSP-coated gold + proteins at different time points (day 1, 3, 7). Values are normalized over the average intensity value of the polystyrene plate control at day 1. Results are presented as average±SD, with N≥12. Statistical significance was evaluated through two-way ANOVA with * indicating p < 0.05; **, p < 0.01; ***, p <0.001, ****, p<0.0001. Cells attached on DSP+BFet, presentsimilar metabolism on day 1 to the cells on DSP and DSP+BFn, On both day 3 and 7 cells on DSP+BFet display a statistically significant decrease in cellular metabolism with respect to the positive control (DSP+BFn), indicating a significant detrimental impact of fetuin-A on cellular metabolism in early proliferation stages.

Upon attachment (day 1), no statistically significant difference was found between the different samples. The results confirm metabolically active cells after 24 hours, suggesting that their detachment after fixation observed is likely not due to poor attachment. Both on day 3 and day 7, DSP and DSP+BFn perform comparably, while DSP+BSA and DSP+BFet both present significantly lower intensities. Over 7 days, BFet has a statistically significant difference compared to the positive control DSP+BFn, but none compared to the negative control DSP+BSA. A similar study on the effect of covalently-immobilized fetuin-A on cellular adhesion, conducted by Oschatz et al ^25^ on PLLA-co-PEG electrospun nonwovens, demonstrated a similar decrease in metabolic activity of osteoblast cells over 2 days of incubation.

The results of the cell metabolic assay suggest that BFet, when immobilized to create a monolayer on the surface through DSP, hinders cellular metabolism. This could explain the lower averages in cell number and surface coverage detected through immunostaining. It should be noted however that a lower cellular metabolism is just one of the markers of interest in determining the complex phenomena of tissue regeneration and osseointegration, and that no relevant detrimental effect toward cell growth was observed in any of the surfaces modified with fetuin-A.

Interestingly, we could observe that the DSP functionalizer alone, upon initial attachment and over 3 days, sustained cellular metabolic activity to the same extent as DSP+BFn, and equally well to the polystyrene cell culture plate used, acting as a control. At both day 3 and day 7, cells cultured onto the DSP-coated surface, without any protein, presented more metabolically-active cells than both DSP+BSA and DSP+BFet. Such result suggests that the NHS-ester may somehow be interacting with the cells, possibly by creating a covalent bond with the amine groups of the proteins present in cellular membranes^50,51^.

Fetuin-A has been linked to tumor growth and metastasis, with conflicting reports regarding its potential to either promote or inhibit tumor development, depending on the specific organs involved. In the context of studies involving carcinogenic cell lines ^23,52–54^, either used for cancer studies or as models representative of osteoblast cell behavior as in the present study, fetuin-A reportedly increased cell growth and proliferation^55,56^. Few studies, however, have addressed the effect of fetuin-A when bound to surfaces, either covalently or through adsorption. In previous work by Vyner and Amsden^57^, the augmented proliferation of fibroblasts over 14 days on polymer substrates of varying stiffnesses was directly related to the amount of fetuin-A present on the surface following adsorption from serum. This effect of the protein was however only observed in the presence of other proteins from serum, suggesting a synergistic effect between fetuin-A and other proteins and growth factors present in plasma. In this work, non-supplemented media was employed to investigate the adhesive capability of the protein in the absence of other supplements, further indicating that fetuin-A alone may not be able to sustain cell proliferation. The immobilization of fetuin-A on PLLA-co-PEG electrospun nonwovens in the work by Oschatz et al., led to higher mineralization of the substrate in physiological fluid given the protein calcium-binding capabilities, and increased cell attachment and spreading of the osteosarcoma cell line MG-63. Similar to the present investigation, Oschatz et al. also observed a decrease in cellular metabolism over 48 hours.

Further studies are needed to fully clarify the efficacy of fetuin-A as a surface modifier, particularly to investigate what other molecules and proteins contribute to the previously observed physiological effects in solution, such as improved cell attachment and proliferation. In addition, while the Saos2 cell line is representative of osteoblastic response for the factors addressed in the present investigation^58^, differences between physiological and pathological cell line responses may influence observations from simplified systems, such as in vitro studies. Similarly, differences in sequence and structure between human and animal proteins, as in the case of bovine and human fetuin-A^59^, may lead to discrepancies in experimental outcomes. Finally, substrate-specific characteristics, as evidenced by Vyner et al^57^, should be systematically probed to correlate protein structure and surface density with subsequent cell response, thereby enabling the identification of robust trends and structure-function relationships.

## 4. Conclusions

This study demonstrated that fetuin-A, when covalently immobilized to the surface presenting DSP-functionalization, bound with higher mass and a more viscoelastic layer compared to albumin, as shown by comparative SPR and QCM-D. Radiolabeling evidenced a strong retainment of protein following surfactant-assistant rinses, confirming the strong bond created through the surface functionalizer. Following the seeding of osteoblast-like cells (Saos2) on the substrates, samples were stained for nuclei counting and metabolic activity analysis. No data on cell attachment were collected for day 1 to assess the cell adhesive capabilities of the fetuin-A, as the rinsing steps following fixation removed cells from the surfaces, possibly indicating limited adherence even after 24 hours.

Osteoblast-like cells grown on pre-coated fetuin-A surfaces showed comparable cellular proliferation over 7 days relative to the other protein controls. On the other hand, a significant reduction in metabolic activity was observed in the Saos2 cells over 7 days, suggesting a marked effect of the protein on cellular metabolism. In conclusion, while surface immobilization of fetuin-A was previously found by others ^25^ to augment the deposition of hydroxyapatite-like compounds from physiological solution, the current study demonstrates that when the protein alone is used for surface modification, fetuin-A does not directly promote osteoblast-like cell proliferation, but instead hinders metabolic activity of Saos2 cells. Further investigations are required to probe whether these effects are specific to the cell line used, the chemical linker employed and any associated changes in surface physiocochemical properties, and most importantly any potential co-dependency on other proteins or soluble factors present in the environment.

## Acknowledgements

Confocal microscopy and in vitro studies were carried out at the Centre for Advanced Light Microscopy and the Biointerfaces Institute at McMaster University.

Dr. João Bronze de Firmino, Dr. Bryan Lee, Dr. Joseph Deering and Dr. John Andersson are acknowledged for scientific discussions and support.

AFM-IR analysis were conducted at Bruker Corporation by Jin Hee Kim.

K.G. and K.N.S. acknowledge funding support from the NSERC Discovery Grant Program (grants RGPIN-2020-05722, RGPIN-2019-06433, and RGPIN-2022-05258, respectively), Mitacs Accelerate Program and Mitacs Globalink Research Abroad Program (Application Ref. IT27778 and IT39044, respectively), the New Frontiers Research Fund (NFRFE-2023-00272), and K.G. from the Canada Research Chairs Program (Tier II Chair in Microscopy of Biomaterials and Biointerfaces). A.M. acknowledges the funding and support of the Foundation Blanceflor Boncompagni Ludovisi, née Bildt, and Mitacs Accelerate Program.

## Author Contribution

**Alessandra Merlo:** conceptualization, investigation, methodology, data analysis, image analysis, visualization, manuscript – original draft. **Jesper Medin:** investigation, data analysis. **Andreas Dahlin:** supervision, funding. **Kathryn Grandfield:** conceptualization, supervision, funding. **Kyla Sask:** conceptualization, supervision, funding. All authors reviewed and edited the present manuscript.

