## Supporting Information for "Functionalization of Gold Surfaces with Dithiobis(succinimidyl propionate) for Immobilization of Fetuin-A and Assessment of the Attachment and Proliferation of Osteoblast-like Cells"

Table 1S. Integration to Table 1 presenting the full dataset of XPS data (atomic %) at take-off angles of 90° comparing the surface composition of the DSP functionalized and protein-immobilized samples, comprehensive of details on the contaminants found on the surface. Data are presented as mean ± range, N=2.

|  | Gold |  | Gold+DSP |  | Gold+DSP+BFet | |
| --- | --- | --- | --- | --- | --- | --- |
| C (1s) | 33.75 | ± 1.55 | 31.15 | ± 0.15 | 47.35 | ± 0.35 |
| O (1s) | 19.00 | ± 0.50 | 12.65 | ± 0.15 | 18.35 | ± 0.95 |
| Au (4f) | 44.15 | ± 1.85 | 51.70 | ± 0.30 | 26.35 | ± 2.05 |
| S (2p) | 0.00 | ± 0.00 | 0.90 | ± 0.00 | 0.95 | ± 0.15 |
| N (1s) | 0.00 | ± 0.00 | 0.00 | ± 0.00 | 6.00 | ± 0.20 |
| Cu (2p) | 0.10 | ± 0.10 | 0.60 | ± 0.10 | 0.25 | ± 0.05 |
| Sn (3d) | 3.00 | ± 0.00 | 3.00 | ± 0.10 | 1.85 | ± 0.25 |

Table 2S. Comparison of XPS peaks (atomic %) at incidence angles of 0° and 60° comparing the surface composition of the protein-immobilized samples (Gold+DSP+BFet). Data are presented as mean ± range, N=2.

|  | Gold+DSP+BFet (0°) |  | Gold+DSP+BFet (60°) |  |
| --- | --- | --- | --- | --- |
| C (1s) | 47.35 | ± 0.35 | 58.35 | ± 1.45 |
| O (1s) | 18.35 | ± 0.95 | 19.30 | ± 0.60 |
| Au (4f) | 25.35 | ± 1.05 | 11.75 | ± 1.25 |
| S (2p) | 0.95 | ± 0.15 | 0.85 | ± 0.15 |
| N (1s) | 6.00 | ± 0.20 | 7.80 | ± 0.80 |
| Cu (2p) | 0.25 | ± 0.05 | 0.00 | ± 0.00 |
| Sn (3d) | 1.85 | ± 0.25 | 2.05 | ± 0.25 |

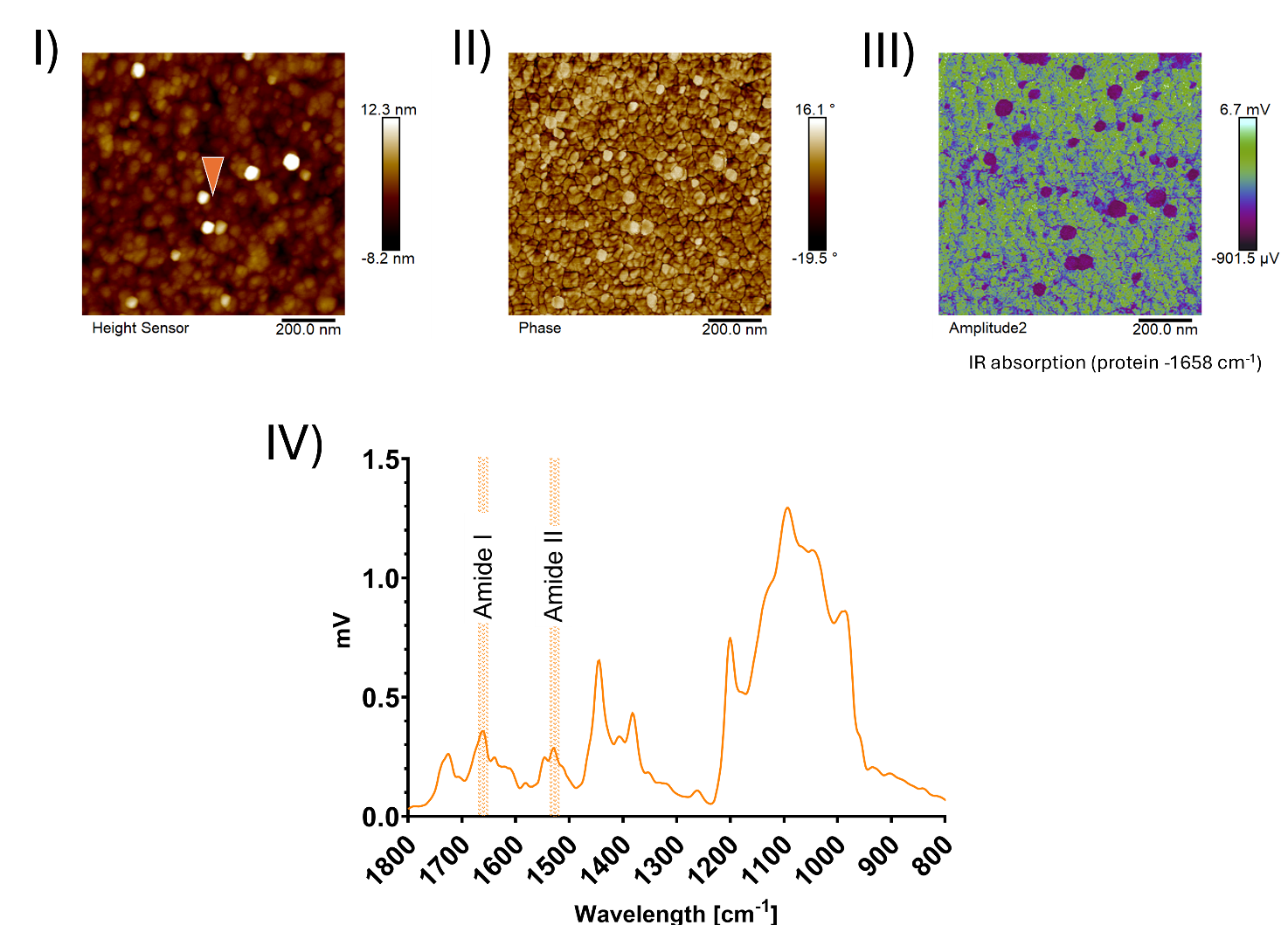

Figure 1S. Atomic Force Microscopy - Infrared Spectroscopy analysis. I) and II) display the morphology and phase lag, respectively, for Gold+DSP+protein. III) displays the cantilever oscillation amplitude from absorption at 1650 cm^-1^, showing the protein distribution on the surface. IV) the IR spectra obtained for Gold+DSP+protein. The areas of the sample where the spectrum was obtained is shown with a downward arrow in the same color of the corresponding spectrum. The amide peaks in G) confirmed the immobilization of proteins.

Both the average peak-to-valley roughness and the phase increased for the Gold+DSP+protein sample, indicating protein binding to the functionalizer, with slightly more viscoelastic characteristics. The absolute value of the phase, however, still indicates a mostly stiff surface, likely because of the gold substrate properties. The presence of white areas in the height measurement could indicate the presence of protein aggregates bound to the surface or some contamination of the sample. The amide I and II peaks (around 1650 and 1550 cm^-1^ respectively) in the Gold+DSP+protein FTIR spectrum confirmed the occurrence of bound protein.

The chemical map from the IR absorption across the surface for the amide I band of the proteins, at a wavelength of 1658 cm^-1^ (Figure 1S, III), demonstrates a relatively spread distribution of the protein on the functionalized surface, but not a full, homogenous, surface coating. The areas that seem to show no indication of protein (shown in dark blue in the image), may be due to contaminants deposited following functionalization and protein immobilization.

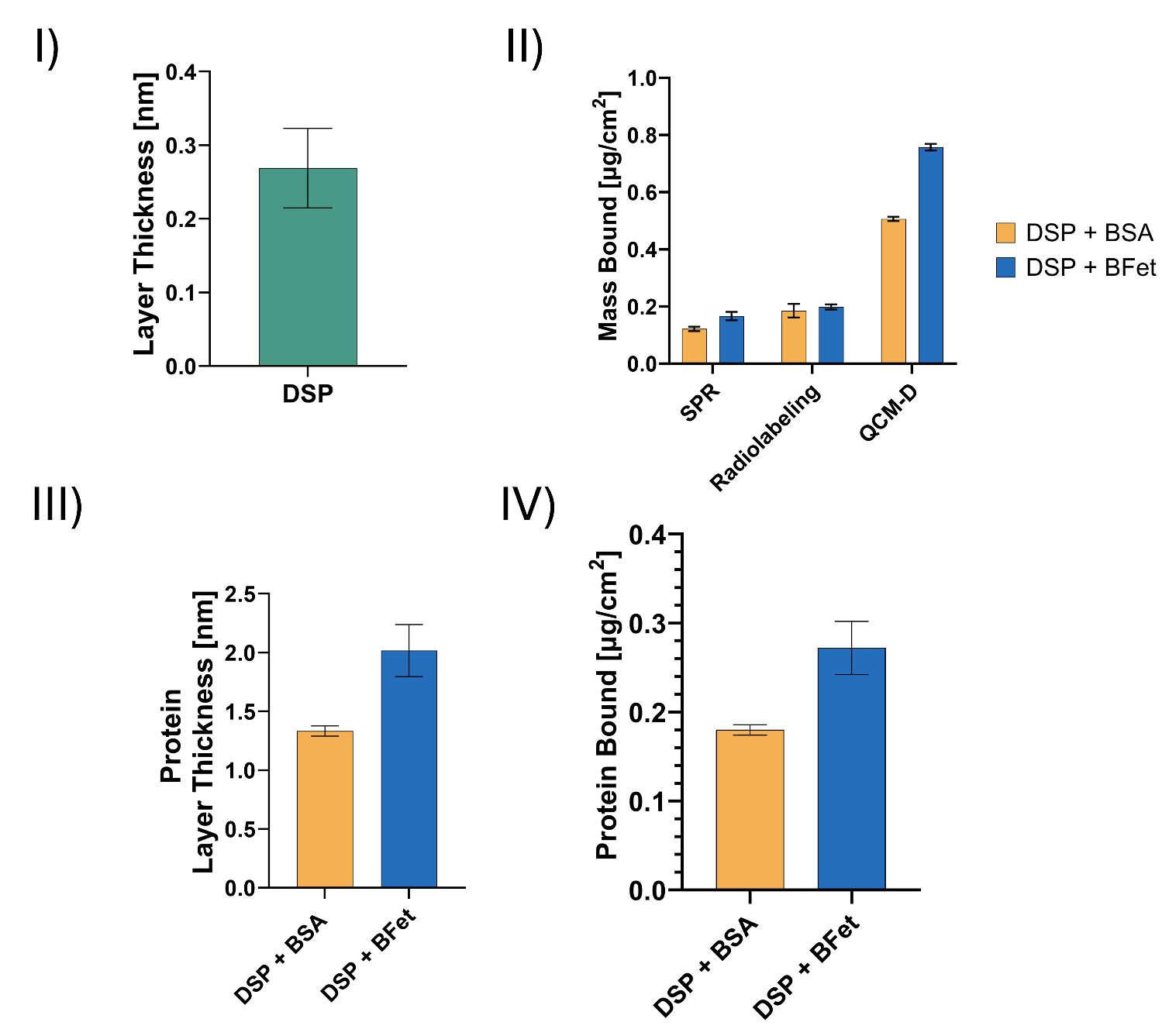

Figure 2S. Histograms represent I) the thickness of the di(thiobis(succinimidyl propionate)) layer deriving from ex-situ functionalization as evaluated from the measured change in SPR angle after 2 h submersion of the samples in 20 mM DSP solution in DMSO solution, followed by water rinsing and nitrogen drying. II) the average mass adsorbed for BSA (yellow) and BFet (blue) at plateau conditions for SPR and QCM-D (40 min injection of a 0.2 mg/mL protein solution in PBS) compared to the mass adsorption measured through radiolabeling (3 h adsorption from 0.2 mg/mL protein solution in PBS under static conditions). III) the thickness of the immobilized protein layer as evaluated from the change in SPR angle after 3 h submersion of the samples in 0.2 mg/mL protein solution in PBS, followed by PBS rinsing and nitrogen drying. IV) the average protein mass adsorbed during ex-situ immobilization, evaluated from the thickness data collected from SPR measurements after 3 h, assuming a 1.35 g/cm^3^ density of the protein layer.

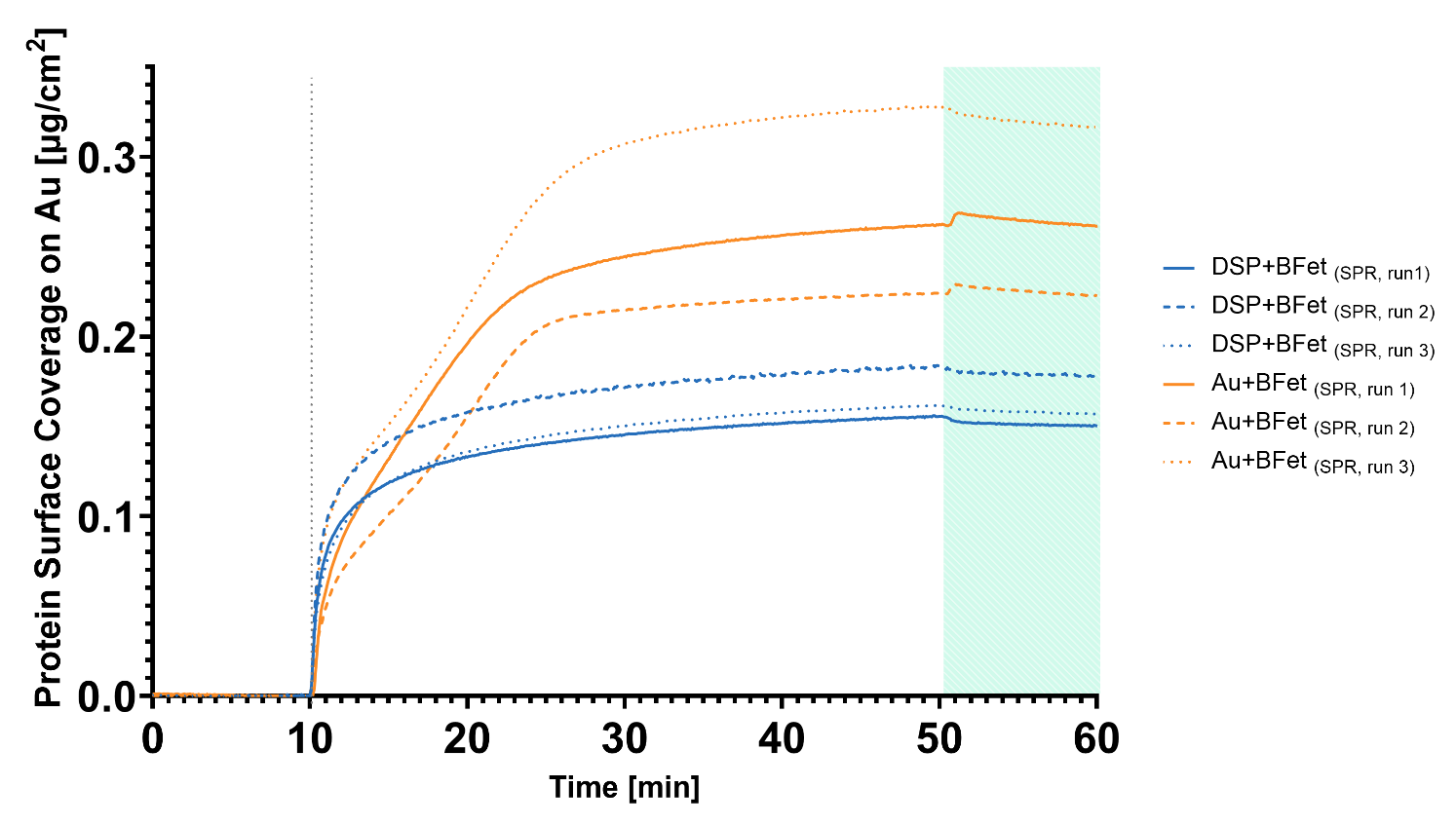

Figure 3S. The graph presents the SPR repeats (N=3) for BFet when adsorbed either on model gold surfaces (in orange) or on DSP-functionalized gold substrates (in blue). The grey dotted line indicates the injection of protein solution (0.2 mg/mL of protein in PBS) following 10 min baseline stabilization with PBS. The green-shaded area indicates instead a PBS-rinsing step following 40 minutes injection, to rinse loosely-bound proteins. BFet surface coverage on gold is higher than DSP-functionalized substrates. Interestingly, while on DSP the binding of the protein follows regular trends, for the adsorption of BFet on bare gold substrates presents more of a linear trend, suggesting a difference in protein interaction with the surface, with more extensive structural re-arrangement in the case of adsorption on bare gold.

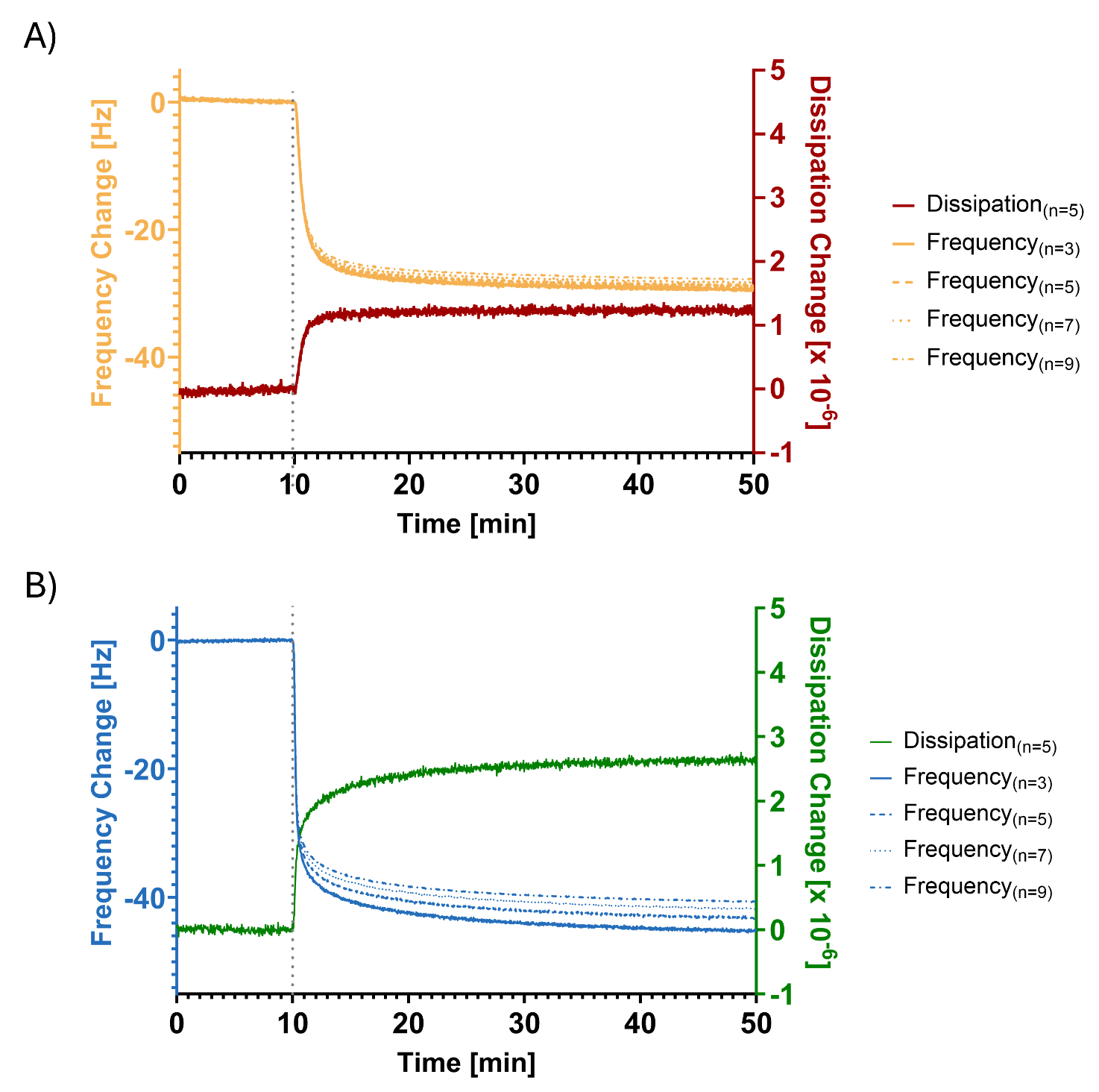

Figure 4S. Representative frequency (3^rd^, 5^th^, 7^th^ and 9^th^ overtone) and dissipation (5^th^ overtone) curves obtained in QCM-D during 40min injections of a 0.2 mg/mL solution of BSA (A) and BFet (B). The grey dotted line represents the injection of protein solution following 10min of baseline stabilization in PBS. The splitting of the frequency overtones for BFet indicates that the BFet layer is more viscoelastic than the BSA layer. However, the limited splitting observed for BSA in the present investigation, when compared to previous results of BSA adsorption on model gold surfaces ^[Merlo et al, bare gold]^,could indicate that the presence of DSP on the surface determines a less rigid composition of the immobilized protein with respect to the adsorbed monolayer.

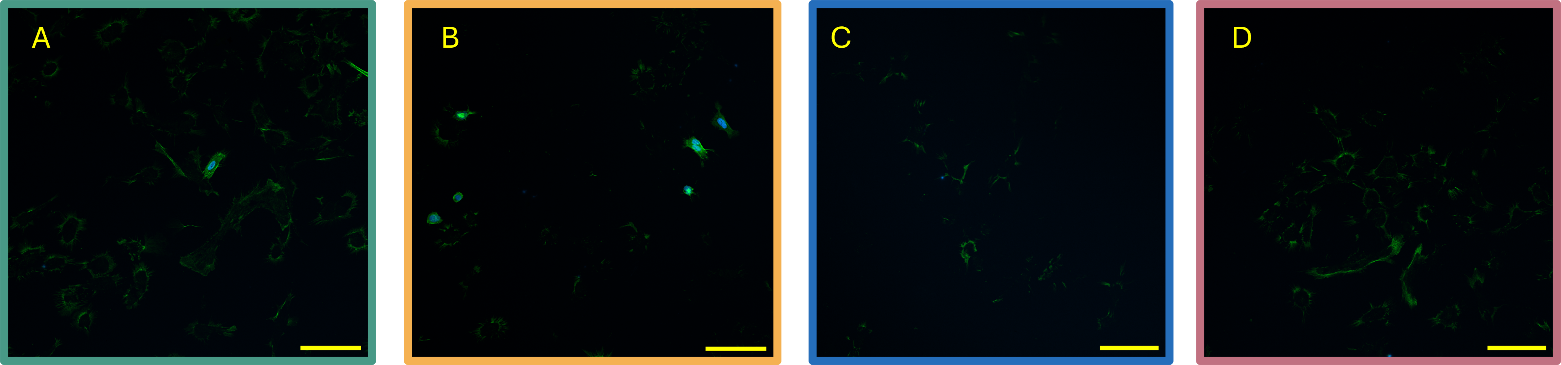

Figure 5S. The fluorescence micrographs show the remnants of cellular debris on the surface following staining and fixation of cells on day 1, for DSP functionalized gold surfaces (A) with BSA (B), BFet (C) or BFn (D). Scale bars represent 100µm.
